# Three-stage processing of category and variation information by entangled interactive mechanisms of peri-occipital and peri-frontal cortices

**DOI:** 10.1101/189811

**Authors:** Hamid Karimi-Rouzbahani

## Abstract

Invariant object recognition, which refers to the ability of precisely and rapidly recognizing objects in the presence of variations, has been a central question in human vision research. The general consensus is that the ventral and dorsal visual streams are the major processing pathways which undertake category and variation encoding in entangled layers. This overlooks the mounting evidence which support the role of peri-frontal areas in category encoding. These recent studies, however, have left open several aspects of visual processing in peri-frontal areas including whether these areas contributed only in active tasks, whether they interacted with peri-occipital areas or processed information independently and differently. To address these concerns, a passive EEG paradigm was designed in which subjects viewed a set of variation-controlled object images. Using multivariate pattern analysis, noticeable category and variation information were observed in occipital, parietal, temporal and prefrontal areas, supporting their contribution to visual processing. Using task specificity indices, phase and Granger causality analyses, three distinct stages of processing were identified which revealed transfer of information between peri-frontal and peri-occipital areas suggesting their parallel and interactive processing of visual information. A brain-plausible computational model supported the possibility of parallel processing mechanisms in peri-occipital and peri-frontal areas. These findings, while advocating previous results on the role of prefrontal areas in object recognition, extend their contribution from active recognition, in which peri-frontal to peri-occipital feedback mechanisms are activated, to the general case of object and variation processing, which is an integral part of visual processing and play role even during passive viewing.

## Introduction

Humans can recognize object categories in fractions of a second with remarkable accuracy (Thorpe et al., 1996; Sofer et al., 2015). This ability seems more outstanding considering that an individual object can produce almost an infinite number of distinct images on the retina as a result of the variations that it undergoes (e.g. size, position, pose, etc.) as well as the variations in the surrounding environment (e.g. background, lighting direction, etc.; Pinto et al., 2008). This has persuaded many researchers to investigate the neural underpinnings of invariant object recognition as a continuously employed sensory-cognitive brain process for everyday vision.

The general consensus is that, the main processing infrastructures of the brain which underlie object category processing are the ventral and the dorsal visual streams (Felleman and Van Essen, 1991; Vaziri-pashkam and Xu, 2017; Pelekanos et al., 2016). The ventral stream starts from V1 and ends up at anterior inferior temporal cortex (IT; DiCarlo et al., 2012; Karimi-Rouzbahani et al., 2017b) and the dorsal stream starts from V1 and continues to parietal and areas in middle temporal cortices (Bullier, 2001; Hebart and Hesselmann, 2012; Freud et al., 2016). In object recognition, linearly-separable representations of objects as well as other information, which are obtained at final layers of the two visual streams, are then sent to the prefrontal cortex for final classification of representations into object categories (DiCarlo and Cox, 2007). However, this view has been challenged by recent studies which have observed category-related information in frontal brain areas, even earlier than they generally appeared in occipital and temporal brain areas after stimulus presentation (Bar et al., 2001; Bar et al., 2006; Kveraga et al., 2007). These latter studies were triggered by a pioneering investigation which reported category encoding in orbitofrontal cortex (OFC; Thorpe et al., 1983). However, systematic investigations are still needed to provide deeper insights into the contribution of frontal brain areas in the processing of object categories and variations as well as possible interactions between anterior and posterior brain areas.

Based on the model proposed by Bar et al. (2006), the orbitofrontal cortex receives low-frequency category-related information from early visual areas (e.g. V1 and V2) through magnocellular pathways (Peyrin et al., 2010; Bullier, 2001), and sends initial guesses about object categories to higher visual areas in inferior temporal cortex (e.g. fusiform gyrus) for more rapid and accurate categorization (Chen et al., 2007; Bar et al., 2001; Bar et al., 2006; Kveraga et al., 2007). In the study performed by Bar et al. (2006), the phase-locking of category-related responses was used to suggest occipital to orbitofrontal flow of information in the window roughly from 80 to 180 ms post-stimulus onset and orbitofrontal to IT cortex flow of information starting at later time windows (after 130 ms). However, as that study investigated average responses, as representative for the amount of information, and did not explicitly measure the flow of category information between the mentioned areas, it remained unknown whether category information was actually transferred between those areas.

A recent study applied multivariate pattern analysis along with Granger causality analysis on magneto encephalographic (MEG) data and investigated the encoding and transfer of category information across peri-frontal and peri-occipital areas (Goddard et al., 2016). That study reappraised the model proposed by Bar et al. (2006), with low‐ and high-spatial resolution image set presented to subjects in a recognition experiment. Nonetheless, the spatiotemporal dynamics of category encoding was drastically different from those reported in Bar et al. (2006), and showed the domination of feed-forward information flow from peri-occipital to peri-frontal areas in early processing time windows (from 0 to around 500 ms post-stimulus) and the domination of feedback flows in the following time windows (from 500 ms to 1200 ms post-stimulus). Authors explained that the observed discrepancy from previous results (Bar et al., 2006) might be explained by the long stimulus presentation time (i.e. 500 ms) in their study which caused the domination of feed-forward information flow in early time windows (Goddard et al., 2016). Therefore, the unbiased spatiotemporal transfer of the category information between peri-occipital and peri-frontal brain areas has still remained to be studied.

Importantly, when investigating the processing of category information in the brain, one needs to always take into account the impact of variations on categorical information and their possible spatiotemporal interactions, as the processing of variation information has been shown to accompany category information in many areas of ventral and dorsal visual pathways (Schwarzlose et al., 2008; Kravitz et al., 2010; Uyar et al., 2016; Hong et al., 2016; Ghodrati et al., 2014; Hung et al., 2005; Li et al., 2009; Kravitz et al., 2013; Carlson et al., 2011). As the compensation of variations has been suggested to be one of the main reasons behind the activation of feedback mechanisms in the brain (Serre et al., 2005; Hupe et al., 1998; Lamme et al., 1998; Wyatte et al., 2012; Karimi-Rouzbahani et al., 2017b), it is interesting to know whether the frontal brain areas also participate in the processing of variations. Except for a few studies, which have addressed the processing of affine variations in the brain (Carlson et al., 2011; Hong et al., 2016), the processing, and more importantly the flow, of variation information in the brain has remained largely overlooked by vision studies.

To address these issues, I developed a whole-brain electroencephalography (EEG) recording experiment in which humans were presented with a set of visual objects which underwent levels of controlled variations. The stimuli were presented very briefly and the task was designed in passive format respectively to allow the appearance of feedback signals in early time windows (Goddard et al., 2016) and to avoid influences from top-down cognitive signals which appear during active recognition (Afraz et al., 2014; Chikkerur et al., 2010; Milner, 1974). Using multivariate pattern analysis (MVPA), I analyzed the spatiotemporal dynamics of category and variation processing in the brain and found the occipitotemporal, parietal and prefrontal areas to be the major areas involved in the processing of categories and variations by entangled mechanisms. I also implemented a recently-proposed version of Granger causality (Goddard et al., 2016) to investigate the dynamical transactions of category and variation information between the peri-occipital and peri-frontal brain areas. Interestingly, I found three distinct stages of information transactions between these areas which support object recognition. These results provide evidence that a set of hard-wired (task-independent) parallel interactive mechanisms within peri-occipital and peri-frontal areas process a combination of category and variation information in distinct stages of processing.

## Methods

### Stimulus set

In order to investigate the processing of object categories and variations in the brain, a four-category object image set with levels of controlled variations was generated. The 3D object models used in the generation of the image set were freely downloaded from (http://tf3dm.com) and rendered using Python commands in the freely available rendering software ‘Blender’ (https://www.blender.com). The image set consisted of four categories of objects including animals, cars, faces and planes, each of which underwent levels of variations in size, position, pose (in-depth rotation) and lighting (Fig. 1A).

**Fig. 1.**
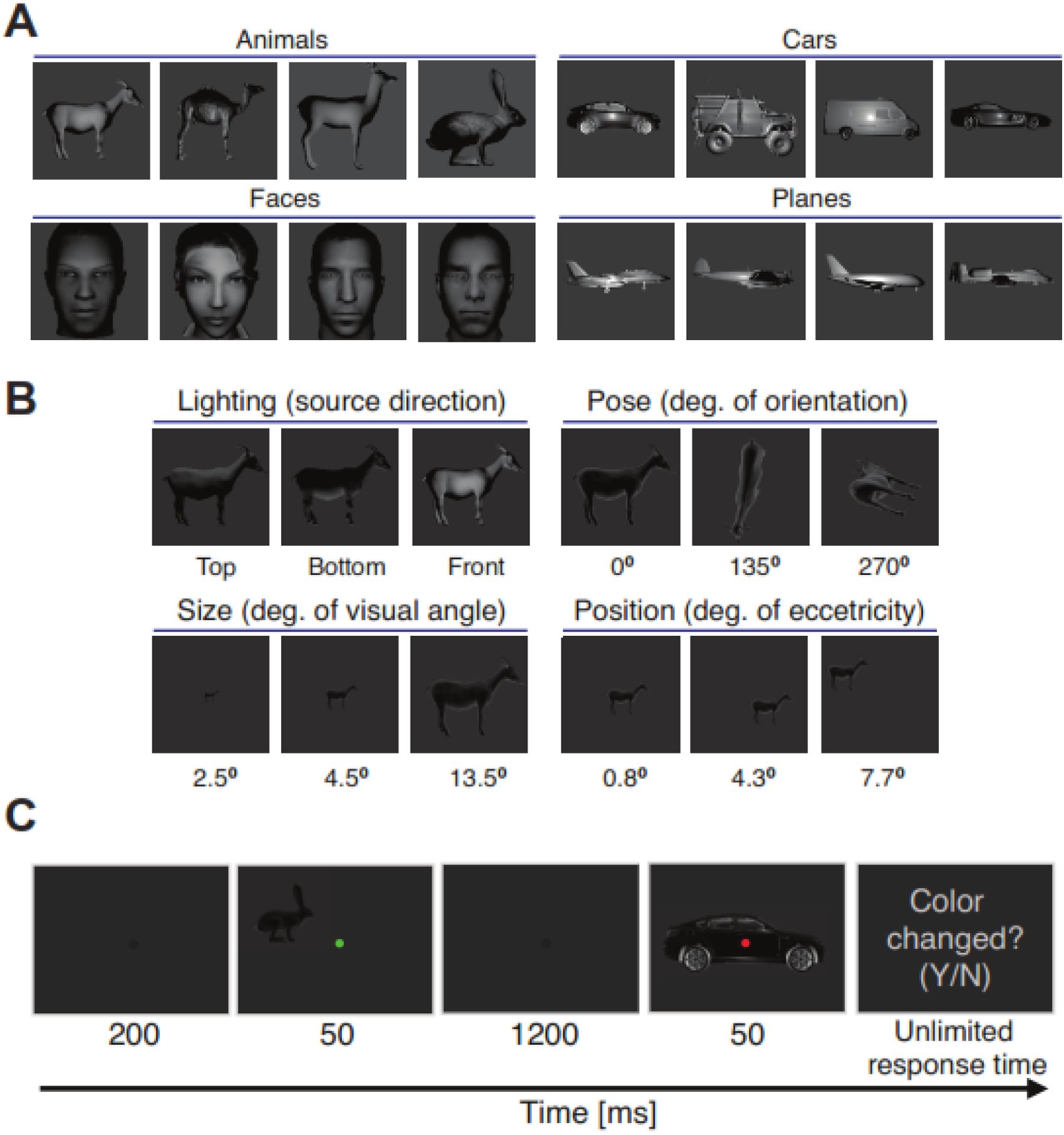
Image set and experimental paradigm. (A) and (B) show the object exemplars within each category (A) and conditions of each variation (B). Images were processed (zoomed in and cropped) for better illustration. Extra information regarding condition are provided below it. (C) EEG recording paradigm with numbers indicating the presentation time of each event.

Each category consisted of four unique exemplars to enhance the generalizability of the image set. In order to cover natural variations of objects, which humans experience in everyday object recognition, I generated images in which objects underwent three levels of variation in size (i.e. 2.5, 4.5 and 13.5 degrees of visual angle) and positioned the objects on three different locations (i.e. with foveal eccentricities of about 0.8, 4.3 and 7.7 degrees of visual angle). I also applied three levels of in-depth rotations on the objects (i.e. 0, 135 and 270 degrees of orientation simultaneously around X, Y and Z Cartesian axes) and illuminated them from three different directions (i.e. top, bottom and front; Fig. 1B). I used a uniform lighting source for all variations except for the lighting conditions. The lighting conditions were selected in a way to present the objects in their most hard-to-recognize everyday conditions, so as to activate all primary and secondary brain mechanisms which are considered to play role in object processing (i.e. including regions in peri-occipital as well as peri-frontal cortices). The final image set consisted of 192 unique images which covered 512 by 512 pixels when presented on the screen during the EEG recording experiment. In order to avoid trivial decoding results, the image set was normalized for equal cross-category and cross-variation average luminance and contrast. Note that the presented images in Fig. 1 are zoomed and chosen from the frontal lighting condition (i.e. 3^rd^ lighting condition) for improved visualization.

### EEG recording paradigm

I used EEG as the brain imaging modality for investigating the temporal and spatial dynamics of object and variation processing in the human brain. EEG could provide signals with high temporal resolution which enabled me to study the highly time-dependent dynamics of object representation in the brain. I designed a passive recording paradigm in which subjects were not supposed to categorize the presented objects, but rather had to attend to category-irrelevant fixation points during the experiment (Fig. 1C). Specifically, at the beginning of each trial, a black fixation spot was presented on the center of the screen for 200 ms after which the first stimulus was presented for 50 ms. Upon the disappearance of the stimulus, an inter-stimulus interval was maintained for 1200 ms before the second stimulus was presented to the subject for 50 ms. The fixation spot remained on the center of the screen throughout the trial, but randomly switched color to either red or green when accompanied the two stimuli. Subject’s task was to decide whether the color of the fixation spot remained the same or changed from the first stimulus to the second (i.e. it changed in 50% of the trials), by pressing one of the two predefined keys on the keyboard after the removal of the second stimulus. Response time was not limited and the subjects had to respond to move to the next trial. The next trial began after either the subjects responded or after 800 ms post-stimulus onset whichever happened later. Ten human subjects (average age 22 years, three females) volunteered for this single-session EEG recording experiment which lasted for about 45 minutes. Informed consent was obtained from each participant. All experimental protocols were approved by the ethical committee of Shahid Rajaee Teacher Training University. All experiments were carried out in accordance with the guidelines of the declaration of Helsinki and the ethical committee of Shahid Rajaee Teacher Training University. Subjects had normal or corrected to normal vision. They seated in a dimmed room 60 cm away and against an Asus VG24QE monitor on which the visual stimuli were presented. Objects’ images covered between 2.5 to 13.5 degrees of visual angle depending on their size conditions. Each unique object image (i.e. 192 images in the image set) was presented to each subject three times in random order (adding up to a total of 576 stimuli). Specifically, each subject was presented with a randomly-ordered presentation of three repetitions of the same 192 images in the dataset, without any consideration of whether the image was to be presented first or second in each trial. The repetition of presentation was aimed at increasing the signal to noise ratio in the analyses. Trials were divided into three blocks with five minutes of resting time between the blocks. Matlab PsychoToolbox (Brainard, 1997) was used for task design, image presentation and response recording. Subjects participated in a short training session before the main experiment on a different image set to get acquainted with the task. Two major considerations were made when designing the paradigm to avoid interfering factors in the results: a) the paradigm was designed in passive format (i.e. subjects performed an irrelevant task and did not actively categorize objects) to avoid the involvement of top-down cognitive processes such as attention and expectation (as they may modulate the dynamics of visual processing in the brain; Pantazatos et al., 2012; Chikkerur et al., 2010; Milner, 1974) and allow only the sensory object processing mechanisms to function; b) images were presented very shortly (i.e. for only 50 ms) to avoid the domination of feed-forward information in the recorded signals to be able to dissociate between feed-forward and feedback flows of information (Karimi-Rouzbahani et al., 2017b; Hong et al., 2016; Goddard et al., 2016).

### Signal preprocessing

A 32-channel eWave32 amplifier was used for signal recording which followed the 10-20 convention of electrode placement on the scalp (see Supplementary Fig. S1B for electrode locations). The amplifier, which was produced by ScienceBeam (http://www.sciencebeam.com/), provided a sampling rate of 1K samples/second which allowed me to investigate the temporal dynamics of information processing in the brain very accurately. The recorded data was taken into Matlab (http://www.mathworks.com/) and all the following analyses were performed on the data in Matlab. I band-pass filtered the recorded signals in the range between 0.5 to 100 Hz to filter-out the DC component as well as the high-frequency noise. I also notch filtered the signals at 50 Hz to block the line noise. In order to remove eye-blink, body and eye movement artifacts, Independent Component Analysis was used as implemented by the EEGLAB toolbox (Delorme and Makeig, 2004). I used ADJUST plugin (Mognon et al., 2011), which statistically evaluated ICA components and suggested the components which represented the mentioned artifacts for removal. I removed the components which were proposed by ADJUST plugin. A total of 154 trials (mean = 2.67%, sd = 1.4%) were removed from the total set of trials pooled across subjects after artifact removal. Signals were then broken into epochs (analysis time windows) which were aligned to the stimulus-onset, in the range from 200 ms pre‐ to 800 ms post-stimulus. Signals were then smoothened using a 5-sample non-overlapping moving average filter to damp the spurious patterns in the signals. No other observable difference was noticed between the results before and after this smoothing.

### Representational analysis

For each subject, after the above preprocessing steps, an ‘X’ data matrix was obtained which was used in the following representational analysis procedure. The ‘X’ matrix was 3-dimensional with 31 rows (representing each electrode’s signal; one electrode was the reference), 201 columns (representing 5 ms time point signals in the range from 200 ms pre‐ to 800 ms post-stimulus onset) and 576 layers (representing the number of trials from subjects who had no removed trials). All the preprocessed signals were used in the analyses, whether they were evoked by the first or the second stimulus in each trial. The ‘X’ matrix included activity values (i.e. voltage values in microvolts) which were obtained from electrodes.

The representational analysis method of the current study has been previously explained in full detail (Cohen and Maunsell, 2009; Karimi-Rouzbahani et al., 2017a). Briefly, I report a decoding index referred to as d’ (i.e. which has also been referred to as sensitivity, decodability, separability, selectivity and discriminability in previous studies (Averbeck and Lee, 2006; Zhaoping, 2014; Grootswagers et al., 2016)) to show how separable the clusters of distinct conditions (either categories or variation conditions) positioned relative to each other in the representational space and how their distributions changed over time (Supplementary Fig. S1A). In the remaining of this section, I explain the decoding procedure using an example for clarification. In order to calculate the separability of the recorded activities between car and face categories at 150 ms post-stimulus time point in the recorded EEG space, I used all 31 rows of the 71^th^ column of the ‘X’ data matrix in layers 1-144 which contained the car data as well as layers 145-288 which contained the face category data. This pair of 144 data points corresponding to the car and face categories provided a pair of 31-dimensional mean vectors which were used for projecting the data points for the two clusters onto the line connecting the two cluster means and reducing the dimensions of the representational space to one. Accordingly, I obtained a pair of 1-dimensional means and a pair of 1-dimensional variance values for the two clusters which were finally used to calculate the decoding value d’ using equation (1):

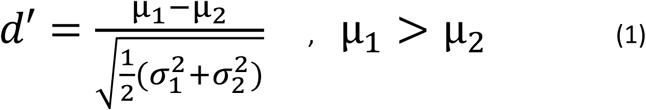

where *μs* and *σ*s are the mean and variance values obtained from the two mentioned clusters (i.e. categories in the above example) and *d’* represents the separability value between the face and car categories. Note that, according to the equation (1), the cluster with a higher mean voltage should be considered cluster ‘one’ to provide positive d’ values, which is actually a distance value and should be always positive. The d’ value was calculated for every time point to obtain the cross-time results depicted throughout the paper (e.g. Figs. 2-4). Moreover, the decoding indices should be calculated between all possible pairs of conditions (e.g. between car and animal, car and plane, etc.). The average decoding value (Fig. 2 and 3, right) for the car category was obtained by averaging all three decoding curves in which the separability of the car category was measured against other categories. To obtain average category decoding results, the decoding indices obtained from the four categories were averaged and reported (Figs. 2 and 3, left). The baseline decoding value (i.e. calculated as the average decoding value within the last 200 ms pre-stimulus window) was subtracted from the corresponding decoding values (within all pre‐ and post-stimulus time points) leading to mainly positive decoding values across decoding curves. The representational analysis procedure was repeated for every subject to obtain individual decoding results, before they were averaged to provide across-subject average results (Figs 2, 3 and 4, shaded error areas represent standard error across subjects). Other decoding methods (i.e. SVM and LDA classifiers) were also tested on the recorded data and led to the same results.

**Fig. 2.**
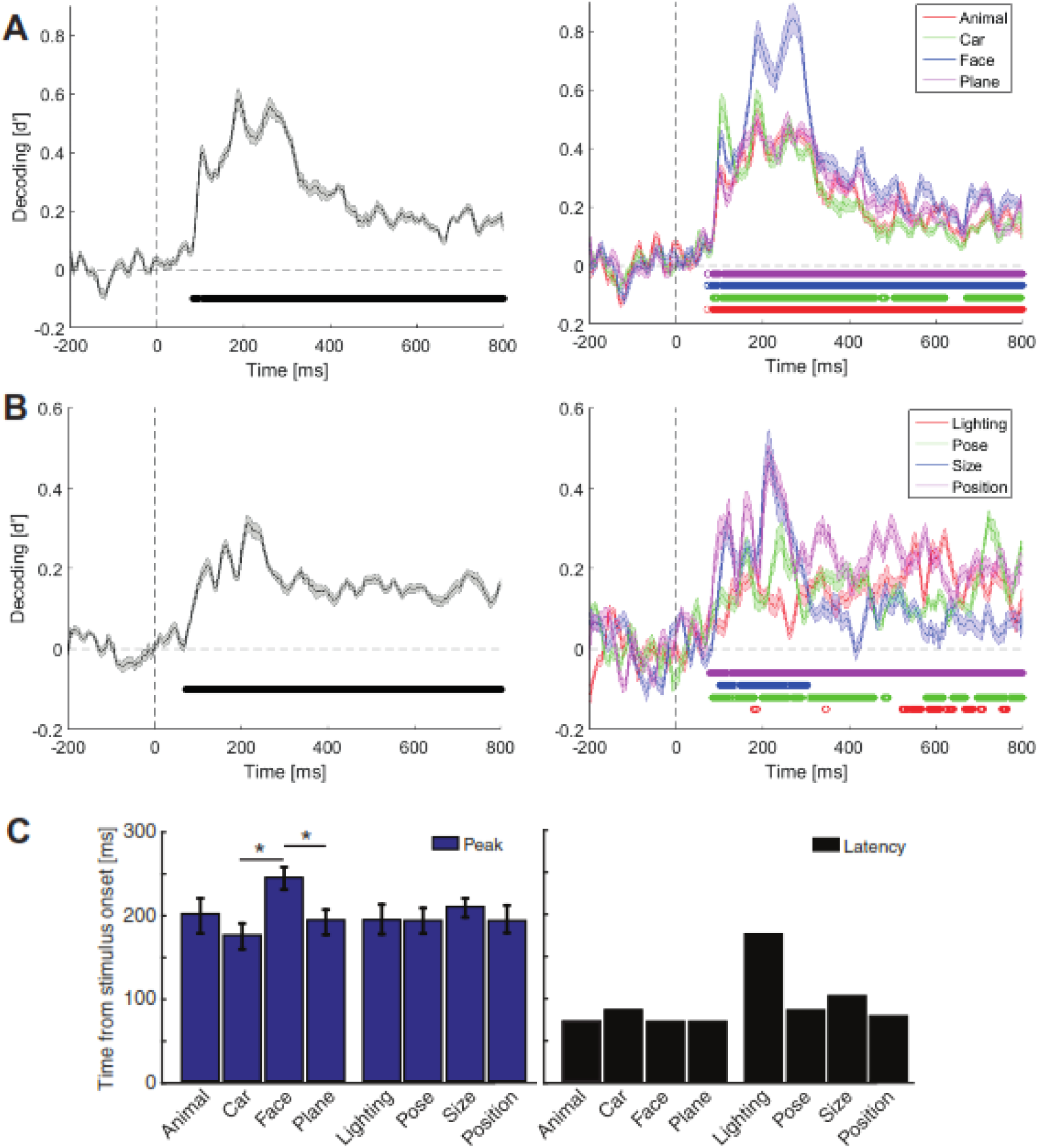
Across-time decoding of categories (A), variations (B) and their temporal statistics (C) for the pooled-condition case. Left columns in (A) and (B) show category‐ and variation-pooled results and right columns show the same results resolved into constituent categories (A) and variations (B). The vertical and horizontal dashed lines indicate respectively the stimulus onset time and the zero decoding value. The circles indicate the time points at which the color-matched decoding curve was significantly above the decoding values averaged in the last 200 ms pre-stimulus (i.e. p < 0.05, Wilcoxon’s signed-rank test). (C) Latency (black) and peak (blue) time points of decoding in category and variation decoding curves. Stars show significant (p < 0.05, Wilcoxon’s signed-rank test) difference between peak time bars. Shaded areas and error bars indicate the SEM across subjects.

**Fig. 3.**
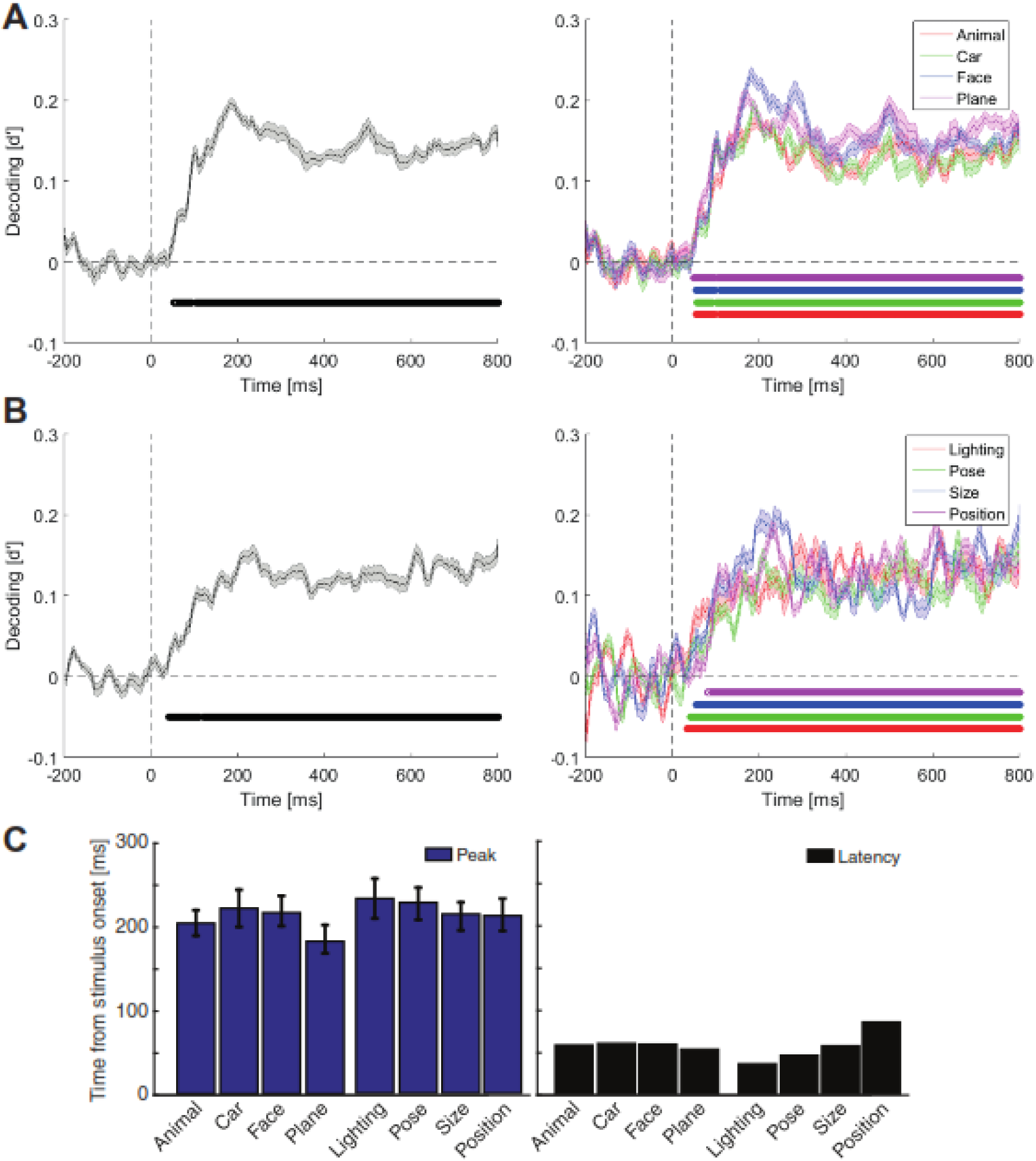
Across-time decoding of categories (A), variations (B) and their temporal statistics (C) in the per-condition case. All the details are the same as in Fig. 2.

**Fig. 4.**
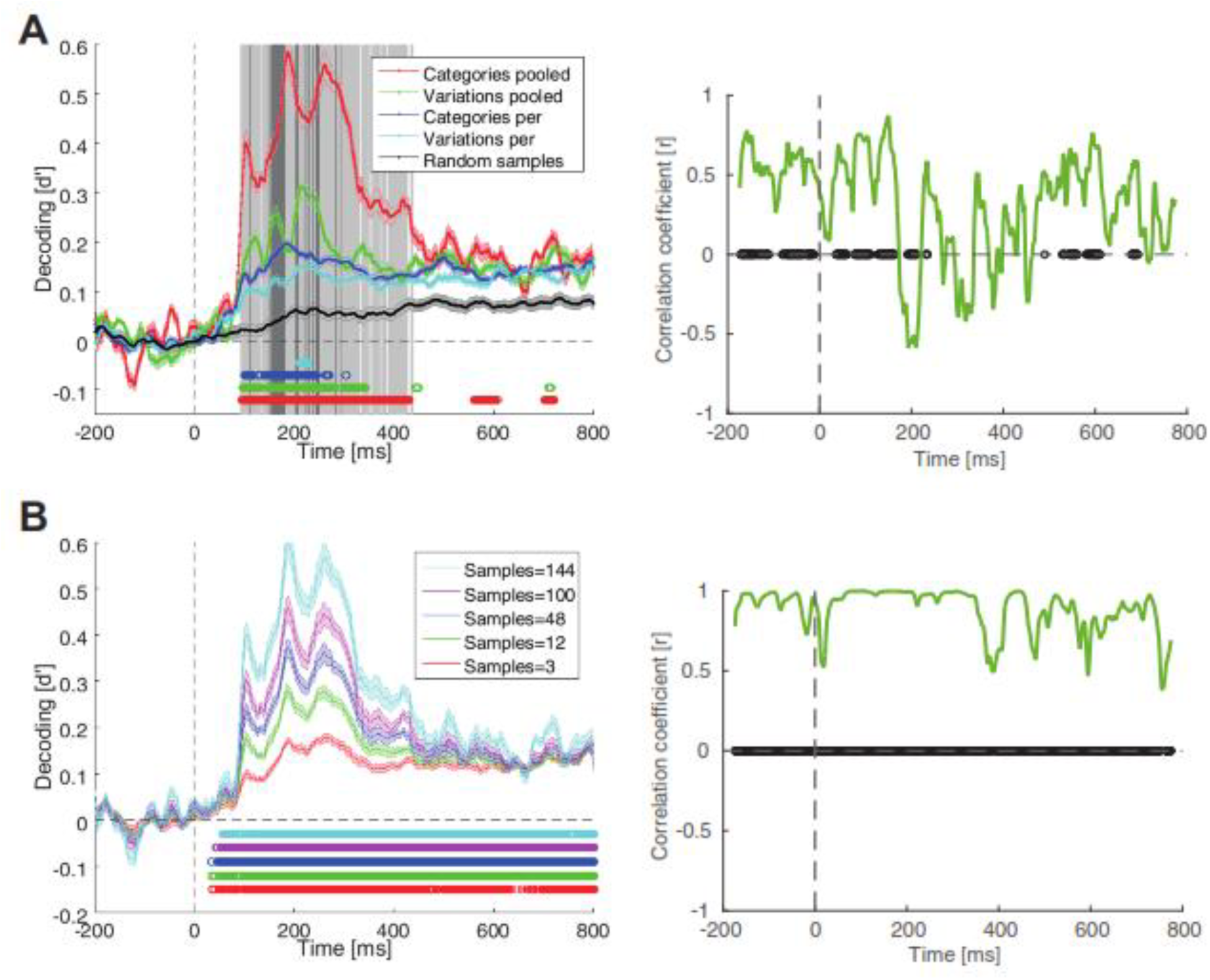
Comparion of across-time decoding of categories and variations in the pooled‐ and per-condition cases. (A) Left, the black decoding curve shows the decoding results of random sub-sampling of the whole stimulus set. The vertical solid light and dark gray lines indicate the time points at which respectively the category and variation decoding curves showed significantly (p < 0.05, Wilcoxon’s signed-rank test) higher pooled-than per-condition values. The colored circles indicate the time points at which the corresponding decoding curve showed a significantly (p < 0.05, Wilcoxon’s signed-rank test) higher value compared to the randomly sub-sampled black curve. Shaded areas indicate the SEM across subjects. (A) Right, the across-time correlation between per-condition cases of category and variation decoding curves. (B) Sub-sampled category decoding curves for different number of samples (left) and the correlation values between the 144 and 3-sample curves (right). The black circles indicate time points of significant correlations (p < 0.05, Pearson linear correlation).

### Granger analysis

In order to investigate the flows of category and variation information between peri-occipital and peri-frontal areas, a recently proposed version of Granger causality analysis was used (Goddard et al., 2016). The logic behind Granger causality is that time series Y might have caused time series Z if Y contains information that facilitates the prediction of future values of Z compared to when considering the information in the past of Z alone. As an example, assume the case of category information moving from posterior to anterior brain areas. In this case, it can be concluded that category information has moved from peri-occipital areas and reached peri-frontal areas if the past representations of peri-frontal alone are not as predictive of the current category representations on the peri-frontal areas as the past representations of peri-frontal plus past representations of peri-occipital are.

First, I needed to have obtained object representations to be able to follow their movement on the scalp. For that purpose, I used the well-known similarity matrices (Kriegeskorte et al., 2008), which provide cross-correlation values between representations. These matrices contain similarity/dissimilarity indices (i.e. indices can be Euclidean distance, correlation coefficient, etc.) which are obtained across brain representations of stimulus pairs (i.e. stimuli can be from the same category in the case of within category analysis, or across variation conditions in the case of variation analysis).

Here, the similarity matrices contained correlation coefficients (obtained from Pearson linear correlation) across pairs of 9-dimensional brain representations (as obtained from nin electrodes) of categories and variations. The dimension of representational space was reduced to nine to separate the peri-frontal and peri-occipital representations. Therefore, two similarity matrices were obtained at each time point; one from the peri-frontal (including F3, F4, F7, F8, FZ, AFZ, FP1, FP2, FPZ) and one from the peri-occipital electrodes (including P3,PF4, P7, P8, PZ, POZ, O1, O2, OZ) (Supplementary Fig. S1B). For example, in order to obtain the similarity matrices of categories, on a single variation condition (e.g. first size condition) for each time point, a 48 by 48 similarity matrix was constructed which included correlation coefficients between all possible pairs of 16 objects (Fig. 1A, each object was presented three times during the experiment). The symmetric sides of the similarity matrices (the top right side which contained values similar to the symmetric bottom left cells) as well as their diagonal axes were excluded from Granger analysis. According to Goddard et al., (2016), partial correlations were used to calculate a simplified version of Granger causality. Equations (2) and (3) provide feed-forward as well as feedback flows of information on the brain on every time point:

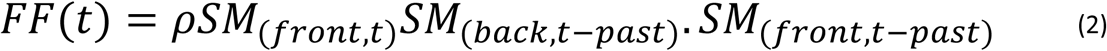

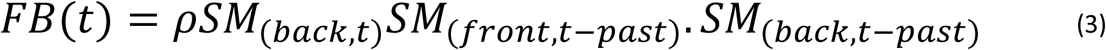

where *SM*_(*loc*,*t*)_ is the similarity matrix obtained from location *loc* at time *t* post-stimulus onset, and *SM*_(*loc,t*-*past*)_ is the similarity matrix which was obtained by averaging the similarity matrices in the window from *t*-130 to *t*-80 ms post-stimulus onset on the same location. The rationale behind this time window was that, it was covered by the range from 72 to 141 ms which was previously shown to reflect the time span for occipital to prefrontal flow of information (Lamme and Roelfsema, 2000). In order to evaluate the significance of the correlation values at each time point, a null distribution of correlation values was generated at each time point by shuffling the elements of the similarity matrices and then using the scrambled matrices in equations (2) and (3). One thousand random correlation values were generated at each time point by repeating the shuffling procedure and calculating the random correlations, against which the true correlation values were assessed for significance. A correlation value was considered significant if it surpassed 95% (i.e. 950) of the random correlation values.

### Computational model

In order to see if a hierarchically organized model of human object processing could provide an explanation for the observed representational mechanisms in the brain, the image set was fed to a recently developed model of human object recognition. The model known as ‘AlexNet’ has been shown to closely replicate object representations obtained from higher visual areas of humans and non-human primates (Khaligh-Razavi and Kriegeskorte, 2014; Cadieu et al., 2014; Karimi-Rouzbahani et al., 2016). I used the model’s Matlab implementation (Vedaldi and Lenc, 2015) which was freely available at (http://www.vlfeat.org/matconvnet/). As the model has been previously explained in many studies (Karimi-Rouzbahani et al., 2017b; Khaligh-Razavi and Kriegskorte, 2014; Cadieu et al., 2014), extra explanations are avoided here. Briefly, the model is an eight-layer convolutional neural network which has been trained on a set of 1000 object categories from the ImageNet Large Scale Visual Categorization (ILSVRC; http://www.image-net.org/; Krizhevsky et al., 2012), including the categories used in the current study, using gradient descent algorithm. The model uses several mathematical operations such as convolution, maximization, normalization and pooling in alternating layers. The first five layers of the model implemented convolutional operations which were followed by three layers of fully-connected units. Since the last layer worked as a 1000-class classifier in previous studies (Vedaldi and Lenc, 2015), I used layers one to seven in the current study. Each output unit at each layer was treated as a representational dimension (i.e. corresponding to EEG channels) and decoding indices were obtained at each layer according to equation (1).

## Results

This study was designed to provide spatiotemporal insight into object category and variation processing at the whole-brain scale. For that purpose, ten human subjects participated in a passive EEG recording paradigm in which they reported if the fixation spot which accompanied the presented objects were the same color or different from the first to the second stimulus in each trial. Subjects performed the task with significantly above-chance accuracy (mean = 95.75%, std = 4.03%, p < 0.001, Wilcoxon’s signed-rank test) and in reasonable time (mean response time = 729 ms, std = 97 ms) meaning that they were alert and attentive to the task during the experiment.

## Temporal dynamics of category and variation processing

To investigate the temporal dynamics of category and variation processing in the brain, I calculated decoding indices (referred to as information in the text) across categories and variation conditions (Fig. 2A and B). The category information (Fig. 2A, left; averaged across all possible pairs of categories) showed a highly dynamical pattern; it rose to significance at 84 ms, experienced three peaks (with the highest peak at 184 ms) and remained significant until 800 ms post-stimulus onset. In order to evaluate the significance of decoding indices at a post-stimulus time point, I evaluated the vector of post-stimulus decoding indices (including ten decoding values corresponding to ten subjects) against their corresponding values averaged in the last 200 ms pre-stimulus window prior to baseline removing, using Wilcoxon’s signed-rank test. Then the p-values were FDR-corrected (using Matlab *mafdr* function) for multiple comparisons and were reported as significant when p < 0.05. Interestingly, the variation information rose to significance earlier than the category information at 71 ms post-stimulus. Although indicating lower peaks compared to the category information, variation information (Fig. 2B, left, averaged across all four variations) presented a similar pattern showing three peaks with the highest at 214 ms post-stimulus. It remained significant until the last analysis time point at 800 ms.

In order to obtain insight into possible differences between different categories and variations, I resolved the averaged results (Fig. 2A and B, left) into its constituent categories and variations (Fig. 2A and B, right). Although all four categories experienced the same three peaks (Fig. 2A, right), higher decoding values were observed for the car category during the first peak (which was at 105 ms poststimulus) which was dominated by the face information during the second and third peaks (which occurred respectively at 188 ms and 271 ms post-stimulus). It should be noted that the reported category information values were calculated between pairs of categories; therefore, a higher information value for face means that, when comparing its separability from the other three categories, faces positioned more separately compared to how the every other category positioned relative to the other three categories. For detailed across-category information plots see Supplementary Fig. S2A.

While rising to significance at an earlier time point compared to the other categories (at 71 ms), the face category also peaked at a significantly later time (mean = 243 ms) compared to car and plane categories (Fig. 2C, left, p < 0.05, Wilcoxon’s signed rank test). Category decoding curves showed undistinguishable latencies which ranged from 71 to 85 ms post-stimulus (Fig. 2C, right). Latency was defined as the time distance from stimulus onset to the first time point at which the decoding index rose to significance. The appearance of face information in the late peaks of category information was explainable by the N170 component of ERP signals which have often been associated with the processing of faces in the brain (Itier and Taylor, 2004). The dominance of face information in the signals can also be explained in light of two considerations: the existence of specialized processing mechanisms which are proposed to be dedicated to face processing in the brain (Freiwald and Tsao, 2010) as well as faces’ small within-category variability of exemplars compared to other categories (see Supplementary Fig. S5B). The previously suggested precedence of intermediate-level (e.g. face-plane and car-face) to subordinate-level (e.g. car-plane) and superordinate-level (e.g. animal-plane and animal-car) category information was also noticeable in the temporal dynamics of category information (Supplementary Fig. S2A; Dehaqani et al., 2016).

I also evaluated the separability of variation conditions. In other words, in was interesting to know whether all variation conditions were perceived (i.e. processed and differentiated) by the brain or not. Within-variation decoding results which were averaged across all pairs of conditions showed that, while conditions of all variations were decodable from brain signals, the conditions of position and size were more distinguishable than the conditions of pose and lighting (Fig. 2B). It means that information about different object sizes and positions were easier to differentiate compared to the conditions of the other two variations from brain signals. This difference could not be explained by the pixel-space separability indices calculated on the image set. Information values (d’) for lighting, pose, size and position variations were respectively 2.95, 1.63, 1.44 and 2.2 in the pixel space. Therefore, as the separability of variation conditions on the brain signals was different from the original ranking in the pixel space, it seems that, as suggested before (Karimi-Rouzbahani et al., 2017b; Karimi-Rouzbahani et al., 2016; Serre et al., 2005), not all variations were processed similarly and compensated for to the same extent by the brain. This can also be supported by the category decoding indices obtained under each of these variations (Supplementary Fig. S2C), with lighting and position showing respectively the lowest and highest impacts on category processing. Lighting rose to significance much later than the other variations, but none of the variations showed a significantly different peak time (Fig. 2C, left).

A previous study named the non-categorical information (i.e. variations), which accompanied the objects as the ‘category-orthogonal properties’, pointing to the fact that these information might be processed by mechanisms which are not necessarily involved in the processing of object categories (Hong et al., 2016). However, in the above analyses, when calculating the category information, the data from all variation conditions were considered in the decoding. Moreover, when calculating the across-condition information of different variations, the data from different category exemplars were considered in the decoding. I thought that, this might have influenced the above results by allowing the interaction of category and variation information in the analyses, as it might have in previous studies (Isik et al., 2014; Hung et al., 2005). More specifically, in the case of category decoding; each of the observed peaks in the category decoding curves (Fig. 2A) might have been evoked by either the dynamics of category processing or the repositioning of data points caused by the processing of variations in the representational space, which could have led to the enhancement of category information. Therefore, in order to nullify the interaction between category and variation information dynamics, I did the category decoding analyses (i.e. calculation of d’) on every single variation condition (to obtain category information) and did the variation decoding analyses on every category exemplars (to obtain variation information, Fig. 3) before finally averaging them. I will call these new analyses ‘per-condition’ and the above analyses ‘pooled-condition’ in the rest of this paper. Although the decoding curves lacked the large bump of information which happened before 300 ms post-stimulus onset and dominated the later decoding values, compared to the pooled-condition case (compare Figs. 2 and 3), the results repeated many of the observations from the pooled-condition. The average category and variation information (Fig. 3A and B, left column) rose to significance respectively at 53 and 41 ms, experienced three bumps in the first 300 ms and remained significantly positive until 800 ms post-stimulus. The dominance of face category decoding and size variation could also be observed. Neither the category nor the variation information showed significantly different peak times across their constituent conditions (Fig. 3C, left). The ranking of category pairs’ information remained almost intact (compare Supplementary Fig. S2A and B) and lighting still provided the least amount of influence on category information (compare Supplementary Fig. S2C and D). See also Supplementary Fig. S3A for the decoding results within each variation in the per-condition case.

## In-phase and anti-phase spans of category and variation processing

As many previous studies have investigated the decodability of category information in the brain (Isik et al., 2014; Carlson et al., 2011; Kaneshiro et al., 2015), two goals were pursued in this study: to investigate the spatiotemporal dynamics of variation processing, and to assess the spatiotemporal interaction between variation and category processing in the brain. ‐

In order to address these issues, I had to first choose either the per‐ or pooled-condition decoding results for the following analyses. Qualitative comparison between per‐ and pooled-condition results (Figs. 2 and 3), done in the previous section, supported their main difference in the early time windows. In order to provide a more accurate insight into the temporal pattern of category and variation processing in per‐ and pooled-condition cases, I provided all average results on a single plot (Fig. 4A, left). As obvious from the curves, the three bumps of information occurred at around the same time in the decoding curves of per‐ and pooled-condition cases of category decoding. This proposes that, as the per-condition decoding curve was obtained on single variation conditions (therefore not influenced by other variation conditions) and showed the same three bumps, the information on the pooled-condition category curves were majorly driven by category information rather than being influenced by variation-related dynamics. After investigating the per‐ and pooled-condition curves of variation decoding, the same conclusion could be made for variation information, supporting minor influence of category information on variation processing. In order to highlight the major window in which the per‐ and pooled-condition cases differed, I indicated the time points at which the information values were significantly higher in the pooled-compared to the per-condition results using light and dark gray vertical lines respectively for the category and variation information. Accordingly, the first and the last time points at which category (and variation) information were significantly higher in the pooled-compared to per-condition were respectively at 92 ms (and 112 ms) and 438 ms (and 294 ms), which were on par with the reported window of sensory visual processing in the human brain (Kaneshiro et al., 2015; Liu et al., 2009) suggesting that the higher number of samples in the pooled compared to the per-condition case in decoding, affected the window of sensory visual processing rather than the later windows of processing which are generally associated with higher order cognitive processes. Nonetheless, I used the per-condition cases to obtain the results provided in the following sections (all results after Fig. 5A) of the paper to avoid unnoticed interactions between category and variation processing.

**Fig. 5.**
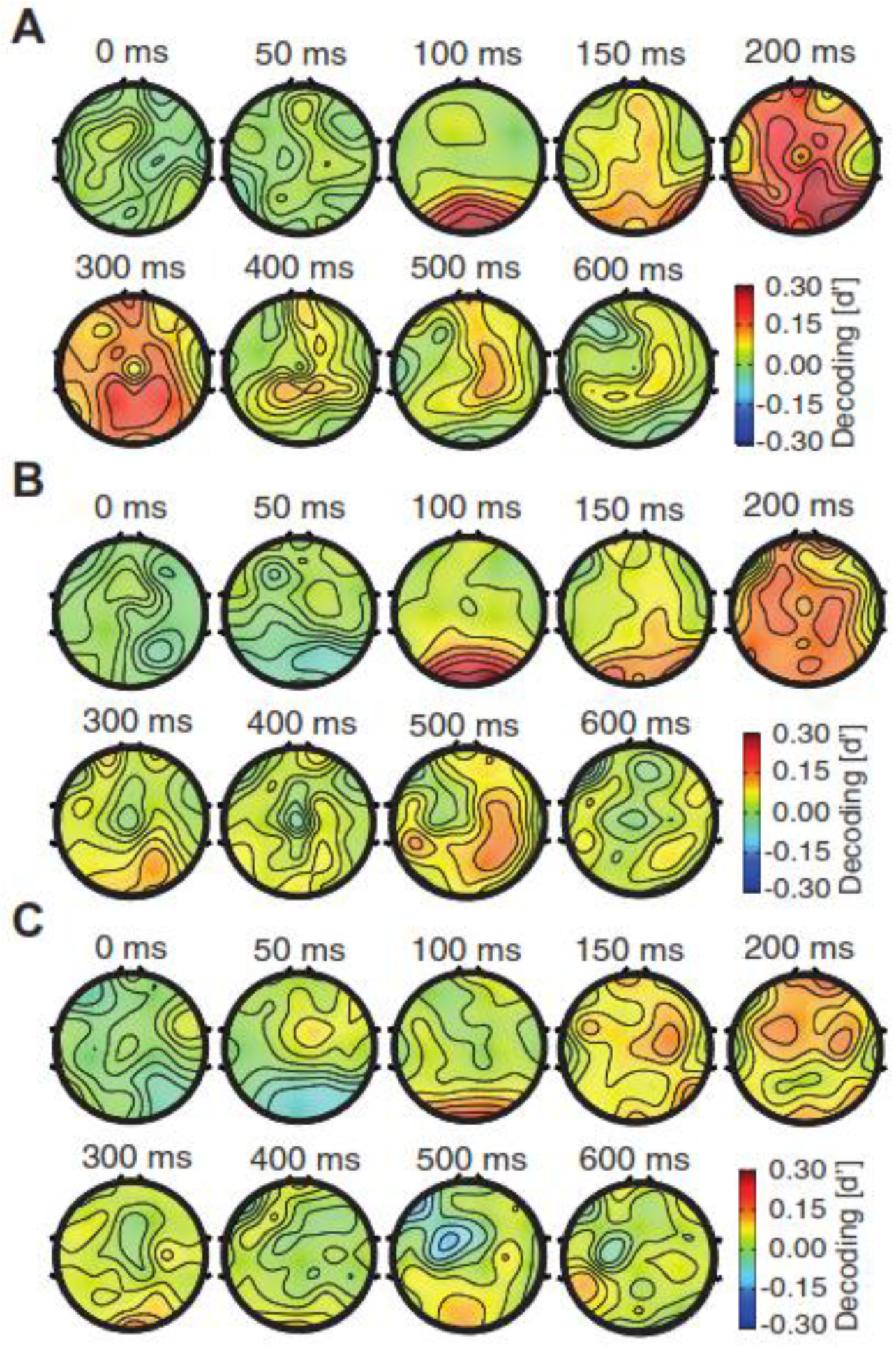
Scalp decoding maps for categories and variations. These maps were generated by measuring the decoding indices for each electrode separately and finally using their superposition on the scalp. The decoding values between the electrodes were calculated by interpolation as implemented in EEGLAB. (A) Pooled-condition category decoding maps. (B) and (C) Per-condition category and variation decoding maps at specific time points. The reported decoding values are averaged in the span from ‐25 m to +25 ms relative to the indicated time points.

Before using the per-condition decoding curves in the following analyses, I had to determine if the calculated decoding indices were significantly above a chance decoding curve which could be obtained from the randomly labeled dataset. This could determine the level of baseline category-unrelated information in the dataset which might have been added to the reported decoding curves. Therefore, I assessed the four mentioned decoding curves against a decoding curve obtained from 144-sample randomly chosen stimulus sets (Fig. 4A, left, black decoding curve). It can be observed that the four curves (even the per-condition curves of categories and variations which consisted of much fewer stimuli) showed significantly above-chance decoding values (as indicated by color circles). The level of significance was very hard to reach for the per-condition variation decoding curve as it included only three data points within each data cluster in the decoding compared to the random sub-sample which had 144 data points within each cluster. In order to generate the mentioned random decoding curve, I randomly selected a subset of 144 stimuli from the whole set of 576 stimuli for each subject, disregarding the category and variation labels of the chosen stimuli, and repeated the random decoding 1000 times before averaging them for each subject data. Together, these results support that the decoding curves, even in the case of per-condition decoding, contained category information which significantly surpassed the information in any randomly chosen subsample of the data which could have information contributing to the reported category and variation information.

An examination of the temporal dynamics of category and variation processing curves (in Fig. 4A, left) suggested that these curves did not follow the same temporal pattern (e.g. compare their peaks and valleys which occurred at around 200 ms post-stimulus). In order to provide a quantitative comparison, I evaluated the across-time correlation between category and variation information curves in their per-condition cases (no noticeable difference was observed when considering pooled-condition cases). Specifically, I calculated the correlation coefficient (Pearson linear correlation) between the time series within the same 50 ms sliding time window of category and variation decoding curves. Correlations were considered significant if their *p*-value was smaller than 0.05 and indicated the corresponding significant time point on the time axis by black circles. Interestingly, while showing a positive value in most other time points, the correlation coefficient curve experienced several systematically negative spans in the window from 173 to 464 ms post-stimulus. Significantly negative correlation coefficients were observed during 192 to 212 ms post-stimulus window. These results suggested three stages of visual processing in the temporal pattern of category/variation processing in the brain: a first stage which started at the stimulus onset and ended at around 170 ms in which information about category and variation was processed in an in-phase manner; a second stage which started at around 173 ms and ended at around 450 ms in which categories and variations underwent several anti-phase processing time spans and a third stage which started at around 470 ms and continued until the end of visual processing with inphase processing of categories and variations. This suggestion will be supported by further analyses in the following sections.

I suspected that the observed difference between the phases of category and variation processing might have been caused by the difference in the number of samples (i.e. number of presented stimuli) considered when comparing category and variation processing. The number of presented stimuli were 144 (and 48) for the pooled and 12 (and 3) for the per-condition cases of category (and variation) decoding. To rule out this possibility, I down-sampled the stimulus set used in the decoding of category data clusters from 144 to 100, 48, 12 and 3 (Fig. 4B, left) and re-calculated the correlations between all possible pairs of subsets, but no negative correlation was observed at any time point (i.e. the sampling procedure was repeated 100 times and the results were averaged before being compared to the true 144-sample decoding curve). In other words, I used a subset of stimuli in this decoding analysis. Results of correlations between the 144- and 3-sample subset, as the most distant cases, are shown in Fig. 4B, right. Therefore, the time-dependent phasic dynamics between category and variation processing seems to be inherent in the brain, and not an effect of the number of samples used in the analyses.

## Spatial dynamics of category and variation processing

In order to evaluate the contribution of different brain regions to the processing of categories and variations, I calculated the decoding indices on the scalp (Fig. 5). Scalp information topographies were important as they could provide insight into the brain regions which were most influential in category and variation information processing. For that purpose, as opposed to the above results which were obtained from all of the 31 scalp electrodes, here I report single-channel decoding indices on time-specific scalp maps (Kaneshiro et al., 2015). In other words, instead of in 31-dimensional space, the representations were evaluated in one-dimensional space. Please note that the decoding indices were interpolated to find decoding values between electrode locations using EEGLAB. Fig. 5A shows the pooled-condition category decoding results on the scalp, which has been the most common type of category information reported previously, which includes both category and variations (Kaneshiro et al., 2015). The amplitudes of decoding values are lower here compared to those reported for the 31dimensional space (Figs. 2 to 4), as a result of dimension reduction in the representational space from 31 to one. The reported decoding values are the average of the decoding values obtained in the time-window from 25 ms before to 25 ms after the indicated time instances. Above-baseline category information was observed in the 50 ms as well as 100 ms windows at AFZ and FZ electrodes with significantly (p < 0.05, Wilcoxon’s signed-rank test) above-baseline information at 100 ms in the whole posterior brain (P3, P4, P7, P8, PZ, POZ, O1, O2 and OZ). In the 150 ms and 200 ms windows, significant category information was observed on occipitotemporal (O1, O2, P7 and P8), parieto-central (P3, P4, POZ, CP1 and CP2) and frontal (F3, F4, FZ, AFZ, FPZ, FP1 and FP2) areas. At 300 ms, parietal (P3, P4, POZ and PZ), central (C4) and frontal areas (FC2, FZ, F4, AFZ and FPZ) showed category information which lost their values in the following time windows (i.e. 400, 500 and 600 ms). Together, these patterns of distribution repeated many previous observations, which have reported the involvement of occipital, occipitotemporal, parietal as well as frontal areas, in category processing in similar temporal dynamics (Hung et al., 2005; Thorpe et al., 1996; Kaneshiro et al., 2015). However, as mentioned earlier, the reported category information may have been influenced by the information from variations as explained earlier. Therefore, I also provided the per-condition category (Fig. 5B) and variation (Fig. 5C) processing scalp maps. Although noisier here compared to the pooled-condition results, the three initial windows (i.e. 0, 50 and 100 ms) of both categories and variations repeated the pooled results. While the category information was more concentrated on occipital electrodes (O1, O2, OZ and POZ), variation information was found on frontal (FC2, FC5 and FPZ) areas in the 150 and 200 ms windows. Parietal (P3, PZ and POZ) information was higher for categories compared to variations in the 200 ms window. In the following windows (300-600 ms), category information was observed on both parietal and frontal areas, while variation information was processed dominantly on occipital and frontal regions. These results which provided separated category and variation information on the scalp, showed evidence supporting both spatiotemporally shared (compare the information in the 100 ms window between category and variation processing maps) as well as distinct (compare them in their 200 ms windows) mechanisms involved in the processing of category and category-orthogonal properties (i.e. variations).

It has been previously suggested that all variations do not necessarily use the same set of brain mechanisms for processing in object recognition. In other words, it has been suggested that while size and position are processed by the feed-forward flows of information in the ventral visual stream, variations such as pose and lighting may need top-down feedback signals from higher cognitive areas such as prefrontal cortex to be compensated for during recognition (Karimi-Rouzbahani et al., 2017b; Karimi-Rouzbahani et al., 2016; Serre et al., 2005; Wyatte et al., 2012). To investigate this, I plotted the scalp maps for each variation separately (Fig. 6). In the 50 ms window, information regarding all variations could be found in the centro-frontal (FC1, FC2, F3, F4, F7, F8 and FZ) brain areas significantly (p < 0.05, Wilcoxon’s signed-rank test) more than it could be found on occipital (O1, O2 and OZ) and parietal (PZ and POZ) areas. In the 100 ms window, information regarding all variations could be consistently found on occipital areas (O1, O2 and OZ). In the 150 ms window, while the information regarding lighting and pose could be observed mainly on the centro-frontal areas (CZ, FC1, FC2, F4, F8, FZ and AFZ) and not at occipitotemporal areas (O1, O2 and OZ), information regarding size and position was mainly concentrated on occipitotemporal areas (O1, O2 and OZ). Almost all variations showed a frontal concentration in the 200 ms window with higher values for size variation (which is probably explained by larger stimuli which evoked higher brain responses). Pose conditions evoked previously proposed co-activation of peri-frontal (F3, F4, FZ, AFZ, FP1, FP2 and FPZ) as well as occipitotemporal areas (P3, P4, P7 and P8; Serre et al., 2005), which may suggest the feedback of pose information from PFC to IT cortex. Lighting information consistently covered the occipito-parietal areas in the subsequent windows (from 300-500 ms). During the same windows, pose information was mainly found on temporal as well as frontal areas enhancing the mentioned possibility for their interactions.

**Fig. 6.**
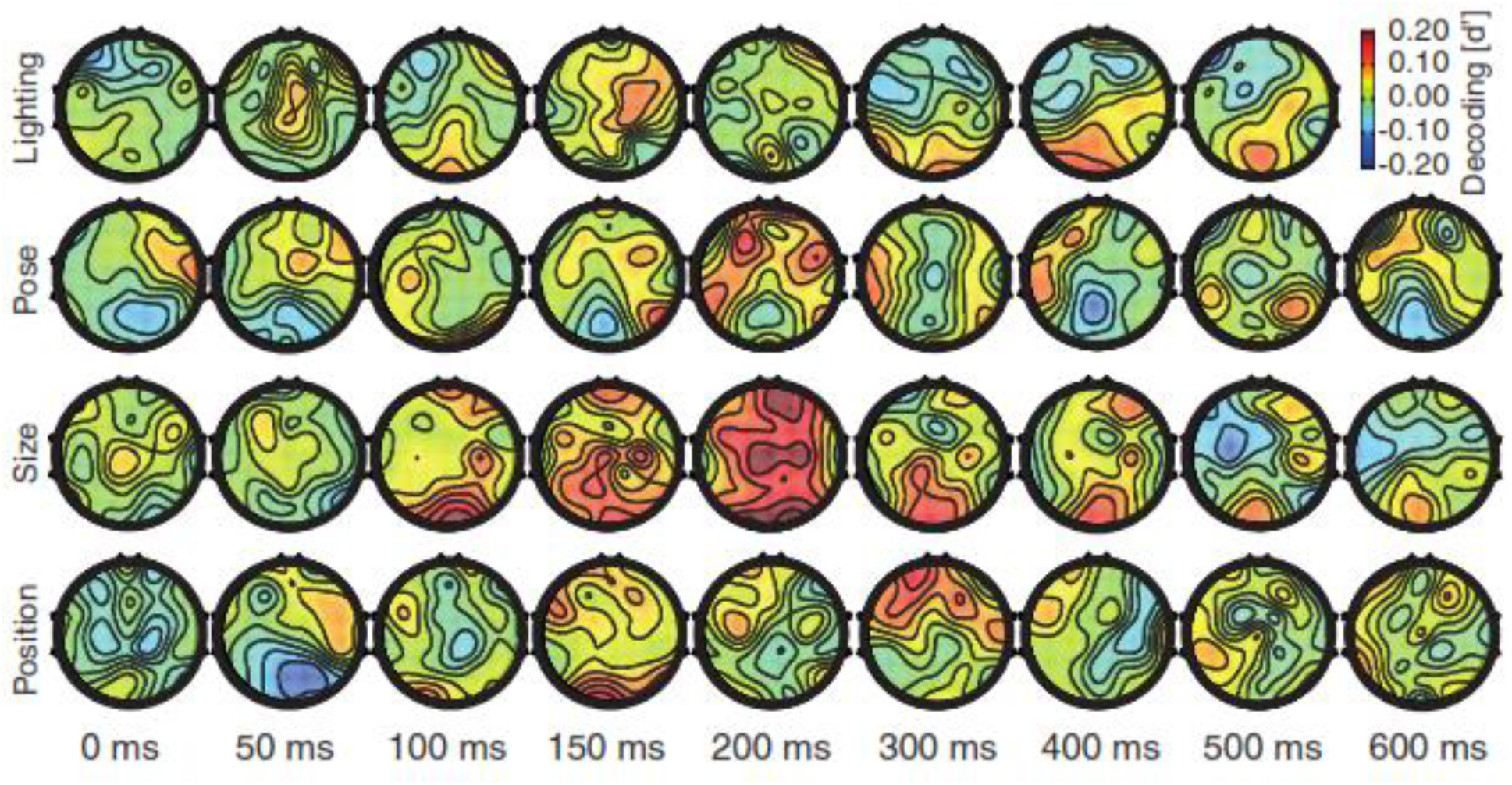
Scalp decoding maps separated for each variation. From top to bottom, decoding maps are provided across conditions of lighting, pose, size and position, respectively.

Interestingly, size information lingered on occipito-parietal areas (O1, O2, OZ and POZ) as well as frontal (F8 and FP2) areas which might be explained by previously suggested frame-transformation in object processing performed in parietal areas (Muthukumaraswamy et al., 2003). As previously proposed by several studies (Karimi-Rouzbahani et al., 2017b; Carlson et al., 2011; Isik et al., 2014)f, position information showed a late appearance in the 300 ms window (see also Fig. 3B, right), and appeared on specific temporal (T7) and frontal areas (F4). By showing that not all variations were processed by the same brain regions, it could be supported that some variations can activate auxiliary mechanisms such as feedback signals from higher cognitive areas of the brain. However, a quantitative evaluation was needed to reveal whether variations and categories are processed by the same brain mechanisms and whether there was any interaction between peri-occipital and peri-frontal areas regarding these processes. In the following sections, these concerns are addressed.

## Task-specificity of the brain to category and variation processing

In order to quantitatively determine whether categories and variations were processed by the same neural structures, I employed a recently-proposed methodology which was developed to evaluate the role of single neurons in the processing of categories and several variations by measuring their selectivity to each of these processes (Hong et al., 2016). I replaced the selectivity indices used in that paper (Hong et al., 2016) by the decoding indices obtained from each electrode here. More specifically, the modified method measured the amount of correlation (Pearson’s linear correlation) at each time point, between the decoding indices found for the vector of 31 electrodes on one task (e.g. processing of animal category) and another task (e.g. processing of size’s first condition). If the set of 31 electrodes provided similar (correlated) patterns of 31-dimensional decoding indices across the first and the second task, the correlation coefficient would be close to unity meaning that the task of animal categorization had evoked the same set of electrodes as did the size’s first condition which implies that these tasks were processed by the same brain areas (and possibly the same mechanisms as the correlations were measured on millisecond time scale). I have provided task specificity matrices (Hong et al., 2016), which reflect color-coded within-and between-task correlation coefficients for every condition at several key time instances (Fig. 7A). The task specificity matrices showed a high-level of task specificity at ‐100 ms prior to stimulus-onset (Fig. 7A). Although instances of confusion between some tasks (e.g. between categorization and position processing, between size and lighting processing, etc.) could be observed, higher values of within-task correlations could be observed compared to between-task correlations. This is not surprising since half of the stimuli had a preceding stimulus whose task specificity was reflected in the pre-stimulus time windows (Fig. 1C). In fact, the decoding indices, which also reflected selectivity (or information), were almost never zero prior to stimulus onset (i.e. that is why I removed the baseline decoding values from the decoding curves); and these non-zero values underlay the observed task specificity in the pre-stimulus span (Fig. 7A). This pattern was also observed in the very late processing time points (e.g. at 328 and 600 ms post-stimulus). Here, however, I concentrated on time instances of significant drops of task specificity which reflected the co-processing (entangling) of tasks in shared brain areas. The processing of tasks have been totally entangled with almost no differentiation between tasks at 50 ms post-stimulus instance as well as at 104 ms. This task overlap faded away at the 200 ms time-instance meaning that distinct brain regions have become involved in the processing of distinct tasks.

**Fig. 7.**
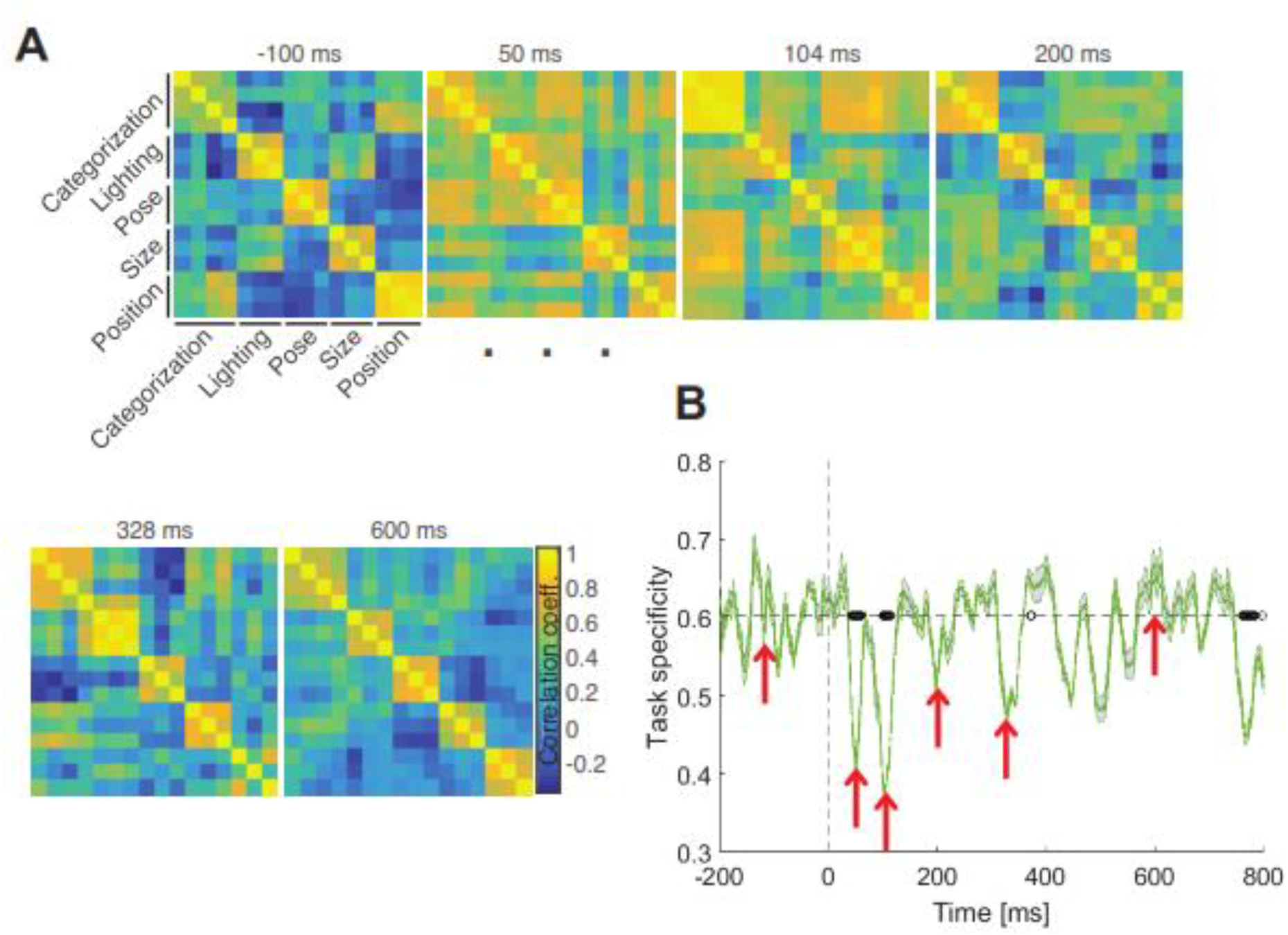
Task-specificity in different brain areas. (A) Task specificity matrices showing, in color codes and at specific time points relative to stimulus onset, the entangling of category and variation processing in the brain. Colors show the amount of correlation (Pearson linear correlation) between decoding indices obtained from whole-brain EEG electrodes in the decoding of specific tasks with higher values showing more similarity (task non-specificity). (B) Across-time task specificity, measured as the difference between the sums of within-task correlations minus the sum of cross-task correlations. Red arrows indicate the time points used in (A). The black circles indicate the time points at which the task specificity index was significantly (i.e. p < 0.05, evaluated using Wilcoxon’s signed-rank test) different from the same index averaged in the last 200 ms window prior to the stimulus onset (before baseline removal). Shaded areas indicate the SEM across subjects.

I also defined and calculated an across-time task specificity index (Fig. 7B). The task specificity index was defined for each task specificity matrix as the average of within-task correlation coefficients minus the average of between-task correlation coefficients. The time instances of the task-specificity matrices, shown in Fig. 7A, are highlighted by red arrows. The task specificity curve revealed its first and second significant (p < 0.05, Wilcoxon’s signed-rank test) declines respectively in the time spans from 43 to 61 ms and from 101 to 113 ms post-stimulus. These significant declines totally matched the occipital processing of category and variation information shown in Figs. 5 and 6. Interestingly, these time spans also highly concurred with the first stage of visual processing (Fig. 4A, right) in which information regarding categories and variations were processed in an in-phase pattern which together support the spatiotemporal co-processing of category and variation information in early visual cortices (Fig. 5). During the second stage of visual processing (from 170 to 450 ms post-stimulus), in which category and variation information were processed in an anti-phase pattern (Fig. 4A, right), distinct tasks were performed by unshared brain mechanisms (Figs. 5 and 6). Together, these results quantitatively suggested that, in the earliest stage of sensory processing, information about categories and variations were processed by the same neural mechanism and in later time windows task-specific brain areas undertook particular tasks.

## Transfer of visual information between peri-occipital and peri-frontal areas

Although the above analyses provided new insights into distinct stages of category and variation processing in the brain, they remained silent on the possible flows of information between brain areas. Recent studies have suggested that specific properties of objects (e.g. low-frequency components of object image) were processed by mechanisms in prefrontal cortex in parallel to the ventral visual stream whose results are transferred from lower (i.e. V1) to higher visual areas such as IT (Bar et al., 2006; Goddard et al., 2016). These studies and other theoretical and experimental investigations, which suggested that some variations may need top-down prefrontal-to-occipital feedback signals for accurate recognition (Karimi-Rouzbahani et al., 2017b; Serre et al., 2005; Tanaka et al., 1996), provided motivation to evaluate the possible transfer of category and variation information between the peri-frontal and peri-occipital areas in object processing. For that purpose, I first separated the information processing at peri-frontal from peri-occipital areas (Fig. 8).

**Fig. 8.**
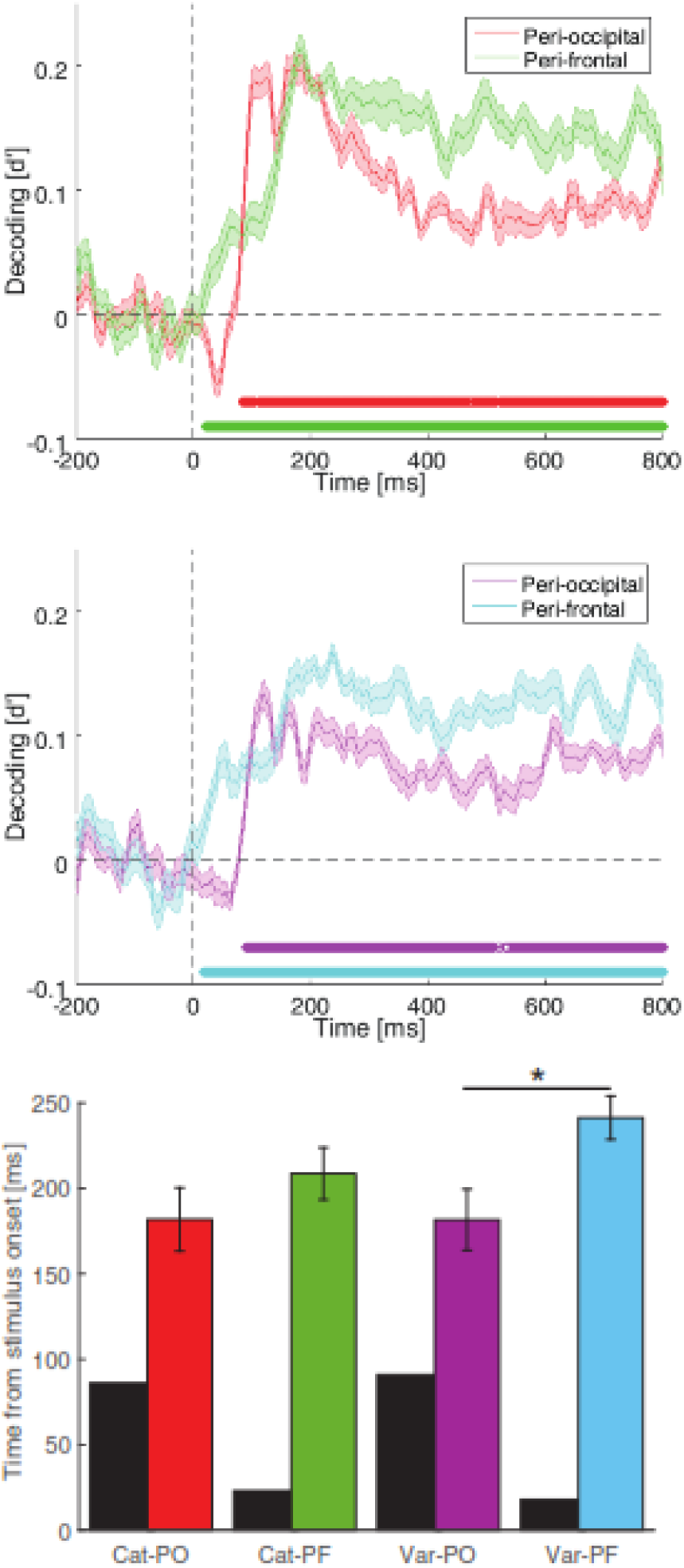
Across-time decoding of categories (top), variations (middle) and their temporal statistics (bottom) in the peri-occipital and peri-frontal areas. The circles indicate the time points at which the color-matched decoding curves were significantly above the decoding values averaged in the last 200 ms pre-stimulus window (i.e. p < 0.05, evaluated using Wilcoxon’s signed-rank test). (C) Latency (black) and peak (colored) time points of decoding in corresponding category and variation decoding curves. Shaded areas and error bars indicate the SEM across subjects. Stars show significant (p < 0.05, Wilcoxon’s signed-rank test) difference between peak time bars. ‘Cat’ and ‘Var’ respectively stand for categories and variations while ‘PO’ and ‘PF’ respectively represent peri-occipital and peri-frontal.

Results showed an earlier rise to significance on peri-frontal information about both categories and variations (respectively at 23 and 18 ms) compared to peri-occipital information (respectively at 86 and 91 ms) (Fig. 8, top and middle). However, information about categories and variations in peri-occipital areas peaked earlier than in peri-frontal areas (Fig. 8, bottom). These results suggest that, in contrast to what might be expected regarding the dominance of category and variation information processing in posterior areas of the brain, the same number of electrodes on the peri-frontal areas can provide even higher amounts of information compared to peri-occipital areas, especially at later stages of processing. This observation was supported by both category and variation information patterns: an earlier rise to significance for peri-frontal information followed by information peaks on peri-occipital and peri-frontal which was followed by higher amounts of information in peri-frontal until the end of analysis time.

Next, I evaluated the spatiotemporal dynamics of information transfer between peri-occipital (including occipital and parietal electrodes of O1, O2, OZ, POZ, P3, P4, P7, P8 and PZ) and peri-frontal (F3, F4, F7, F8, FZ, AFZ, FP1, FP2 and FPZ) areas using a simplified version of Granger causality as suggested previously (Goddard et al., 2016). For that purpose, I first calculated similarity matrices at every time point using the 9-dimentional representational space (i.e. nine electrodes) for object categories and variation conditions.

Then, using partial correlations between representations at time *t* and the average of representations in the span from *t* – 130 to *t* – 80 ms, I investigated the transfer of information between peri-frontal and peri-occipital brain areas (see Methods and (Goddard et al., 2016) for more information). I called the information directions from peri-occipital to peri-frontal as “feed-forward” and from peri-frontal to peri-occipital as “feedback” anatomically and not based on the classical feed-forward and feedback flows of visual information which is dominant in the literature. In other words, rather than the role of peri-frontal areas in higher level cognitive processes such as attention, decision making, etc. (Chikkerur et al., 2010; Milner, 1974), I am investigating the role of specific compartments within that area such as orbitofrontal cortex in providing a parallel processing pathway to sensory object processing (Bar et al., 2001; Bar et al., 2001; Goddard et al., 2016). This is discussed in more detail in Discussions.

Partial correlations between peri-frontal and peri-occipital information showed higher values for category than for variation information (compare red and green curves with cyan and magenta curves in the top panel of Fig. 9A). In order to measure the dominance of information flow on the scalp, I calculated the difference between feed-forward and feedback information (i.e. between partial correlations, Fig. 9A, bottom) and evaluated the significance of the calculated differences against 1000 randomly generated partial correlation values (see Methods). The difference information curves showed highly dynamical patterns switching from feed-forward to feedback at around 150 ms and reversing to feed-forward at around 420 ms for both categories and variations. Results showed that variation and category information led to significant feed-forward flows respectively at 77 and 97 ms and remained significant respectively until 88 and 128 ms post-stimulus. Then, the variation and category information turned into significant feedback flows at 147 and 158, remained significant respectively until 408 and 397 ms, turned into feed-forward flows again respectively at 469 and 448 ms and remained significant until the end of analysis time. Therefore, information regarding categories and variations moved dominantly from peri-occipital areas towards peri-frontal areas in the window from the stimulus onset to 130 ms (first stage) post-stimulus, then back to peri-occipital areas from around 150 ms to 400 ms (second stage) and then again forth to peri-frontal areas from around 450 ms (third stage). The observed stages of category and variation processing confirmed many of the results reported above, as follows. The first stage, which supported feed-forward flows of information from peri-occipital to peri-frontal areas (Fig. 9A, bottom) co-aligned with the in-phase (Fig. 4A) entangled (Fig. 7B) processing of category and variation information.

**Fig. 9.**
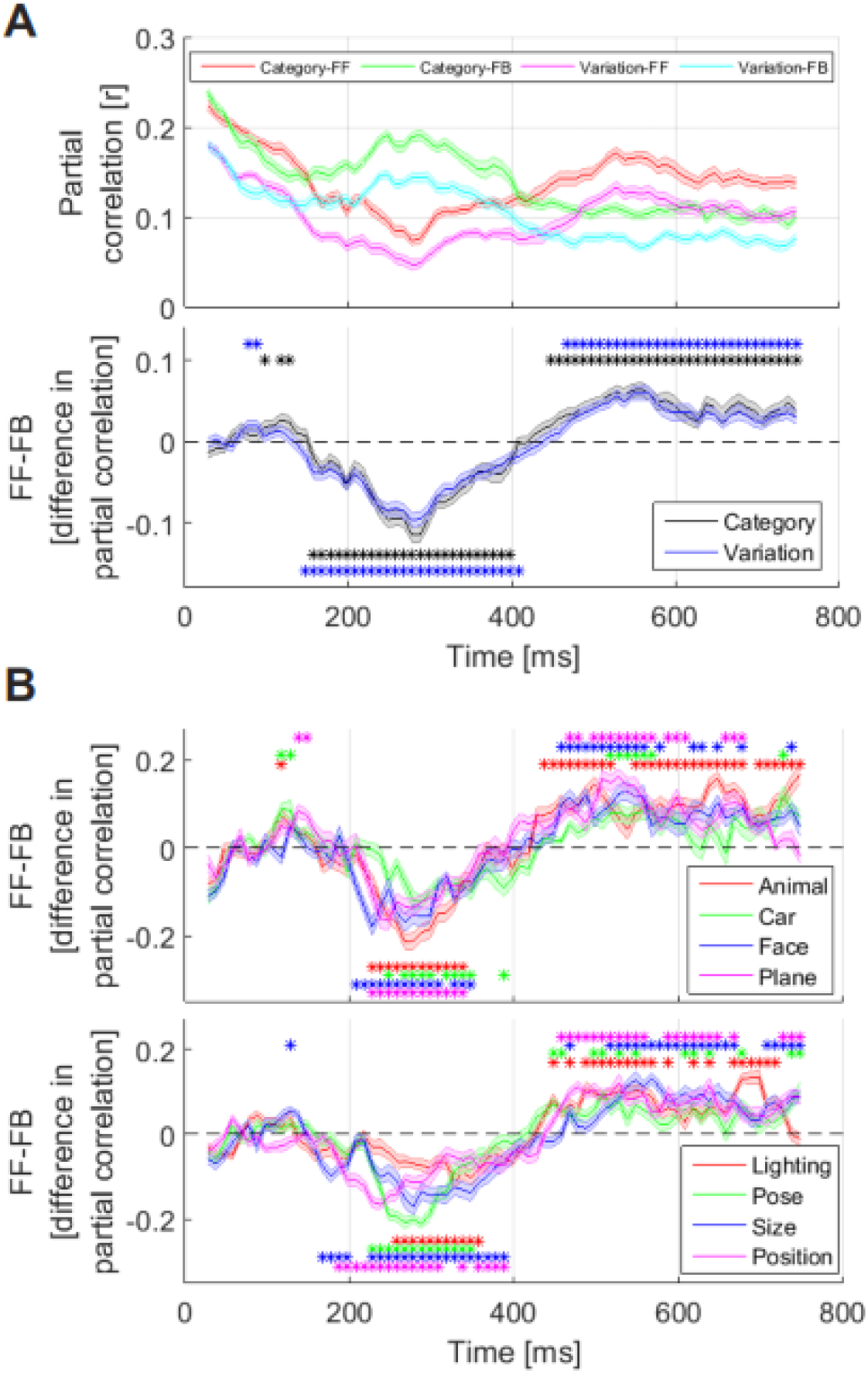
Across-time flows of category and variation information in the brain. (A) Top, partial correlation of representations between peri-occipital and peri-frontal areas. FF and FB refer to the feed-forward (correlation between time *t* representations in peri-occipital and representations during time *t* — 130 to *t* — 80 ms in peri-frontal areas) and feedback information flows, respectively. (A) Bottom, the difference between FF and FB flows of information for categories (black) and variations (blue). Stars indicate the time points at which the flows were significantly higher (p < 0.05, random permutation test) than correlations obtained from a null distribution. (B) The same as (A, Bottom) but for each category (top) and each variation (bottom) with their corresponding significant time points indicated with stars. Shaded areas indicate the SEM across subjects.

The second stage, which concurred with feedback flows of information from peri-frontal to peri-occipital areas (Fig. 9A, bottom), covered the anti-phase (Fig. 4A) task-specific (Fig. 7B) processing of categories and variations. The third stage, which again revealed feed-forward flows of information from peri-occipital to peri-frontal areas (Fig. 9A, bottom), covered the in-phase (Fig. 4A) task-specific (Fig. 7B) processing of categories and variations. The observed temporal dynamics of information flows were also highly consistent with a seminal study which reported feed-forward flow of object information from occipital area (V1) to orbitofrontal cortex at around 80 ms (i.e. the start time of my first stage of processing) followed by the feedback of category information at around 130 ms (i.e. the start time of the second stage of processing of the current study) post-stimulus onset (Bar et al., 2006). To be able to compare the dynamics of information flow across categories and variations, I provided category and variation information flows resolved in their constituent conditions (Fig. 9B). No significant difference was observed between categories (Fig. 9B, top). However, as suggested previously (Karimi-Rouzbahani et al., 2017b), pose seems to have employed a higher level of feedback compared to other variations (Fig. 9B, bottom). More importantly, while category information has provided a higher number of significant feed-forward time points during the first stage, variations have employed a wider span of significant information feedback.

## Comparing the brain’s dynamical behavior with a computational model

A set of computational models of human visual processing have been proposed recently which were able to provide accurate prediction of object representations at final layers of the ventral visual stream (V4 and IT; Khaligh-Razavi and Kriegeskorte et al., 2014; Cadieu et al., 2014; Krizhevsky et al., 2012). One of these models, HMO, which had a hierarchical feed-forward structure, has recently suggested that information regarding both categories and variations were enhanced as object representations passed through layers of the model (Yamins et al., 2014), supporting that a unified structure can process category and variation information by entangled mechanisms. However, as here I supported the existence of parallel pathways for visual object representations, it was interesting to know how one of the most brain-plausible versions of these hierarchical models, known as ‘AlexNet’ (Krizhevsky et al., 2012), would process category and variation information. To that end, I fed the model with the same image set as used in the EEG experiment and measured the decoding indices across categories and variation condtions at the output of every model layer (Fig. 10A). Information about categories and variations increased as object images passed the first layer of the model. Except for lighting which showed a monotonically decreasing information curve, other variations generally experienced information enhancement by going from the first to the fourth layer of the model. After the fourth layer, the variation decoding indices decreased while the category information kept increasing until the last layer of the model. Therefore, two distinct stages of information processing seemed to be at work in the model: one from the first to the fourth model layer and one from the fourth to the last layer. In order to evaluate the relative phase of category an variation decoding in the model, I calculated the correlation of decoding patterns in every three consecutive model layers between category and the average of four variation decoding curves (Fig. 10B), which showed an in-phase followed by an anti-phase processing pattern. These two stages seem to repeat the first and second stages of visual processing obtained from EEG signals (Fig. 9). The third stage of processing, however, was absent from the computational model. This seems to be a result of the model lacking the decision-related mechanisms which were present in the brain and were most probably the destination of information flow during the third stage of visual processing (i.e. PFC). Therefore, the hierarchically-organized feed-forward model of visual processing which was used here seemed to be a brain-plausible representative which implemented the parallel visual pathways (i.e. one going from V1 to orbitofrontal cortex and back to IT and the other directly from V1 to IT) that process visual information prior to the convergence of information at IT cortex.

**Fig. 10.**
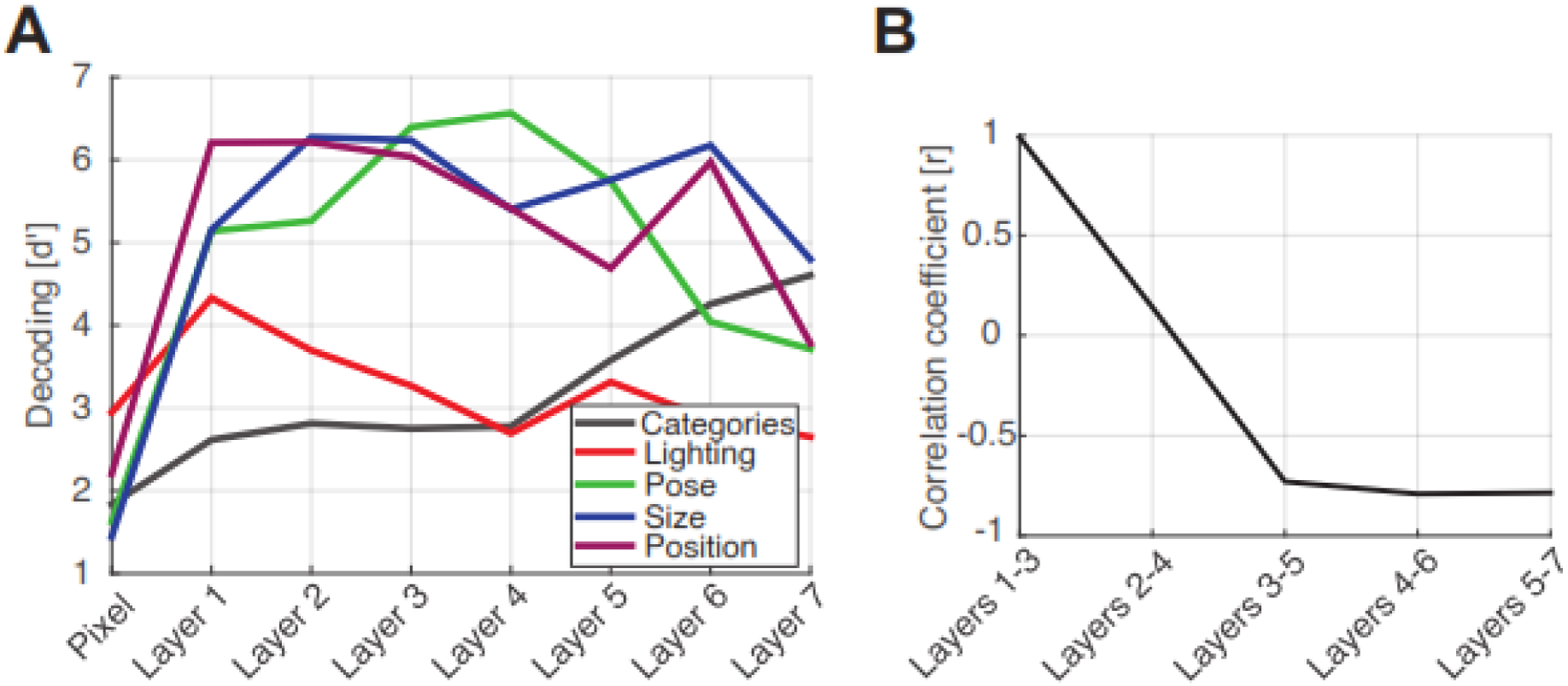
Correlation of the model with brain data. (A) Decoding indices obtained from pixel-space images as well as their representations at the output of every layer of the model for categories (black curve) and variations (color curves). (B) Correlation values between the average of four variation indices and the category index at three consecutive layers of the model (i.e. three data values were used in the calculation of correlations) which showed significantly (p < 0.05, Pearson linear correlation) positive and negative values respectively in the first-half and second-half of the model layers.

The parallel processing structures of the model were probably implemented by different convolutional spatial filters each of which extracted and processed different sub-band frequencies of the visual input which had been inspired by the brain mechanisms for filtering different spatial frequencies (Bar et al., 2006; Goddard et al., 2016).

I also evaluated the spatiotemporal correlation between the category information in the EEG signals, whose results reflected the hierarchical structure of computational models (Supplementary Fig. S4). Results of this analysis confirmed the existence of a hierarchical structure for category processing in the brain, as previously observed for the same model (Cichy et al., 2016). In fact, the results showed that a layer-wise structure could have underlay the observed EEG decoding indices, but does not rule out the possibility of parallel visual processing being at work in peri-frontal and peri-occipital areas. Next, using the representational vectors which were obtained from the last layer of the computational model on an extended version of the current image set (Karimi-Rouzbahani et al., 2017b), the representational dissimilarity matrices also showed the distinctness of face category exemplars from the other categories validating the results explained for Fig. 2A, right (Supplementary Fig. S5A, left). The variation representational dissimilarity matrix also showed the decodeability/distinctness of different levels of variations (size and pose conditions, Supplementary Fig. S5A, right). Finally, using the same extended image set (Karimi-Rouzbahani et al., 2017b) and the representations obtained from the last model layer, I also showed that variations in pose could drastically entangle object representations, whereas lighting had little impact on object representations (Supplementary Fig. S5B and Fig. S2C). This was also reflected in the behavioral object recognition performance of my recent work (Supplementary Fig. S5B, light-colored images) (Karimi-Rouzbahani et al., 2017b). Therefore, although they may divide the visual object processing problem into several sub-problems (e.g. by using sets of convolutional filters with different shape‐ and frequency-based sensitivities) which may not necessarily follow those implemented by the brain (e.g. dividing the problem into low‐ and high-frequency components in frontal and occipital brain areas) (Karimi-Rouzbahani et al., 2017c), the recently developed computational models of human vision can be proper candidates to access primate’s high-level visual representations.

## Discussions

This investigation provides a broad-based survey of the spatiotemporal dynamics of category and variation processing in the human brain. Findings of this study provided several insights. First, the visual processing of category and variation information was shown, for the first time, to be divided into three stages (Figs. 7-9): stage one, which covered the time window before 130 ms post-stimulus during which information about category and variation were processed by (temporally and spatially) entangled mechanisms mainly concentrated in primary visual cortex and were simultaneously (significantly during 80 to 130 ms window) sent to the frontal brain areas (Bar et al., 2001; Bar et al., 2006); stage two, which covered the time window from 150 to 400 ms post-stimulus during which category and variation information, being processed by partially anti-phase distinct mechanisms, were sent back to peri-occipital areas; and stage three, which started at around 450 ms with category and variation information being sent in an in-phase manner to frontal areas possibly for final category-related decisions. Second, this study provided experimental support that information regarding categories as well as variations were processed also in the peri-frontal areas of the brain (Fig. 5). This added evidence to the previously suggested role of frontal areas in the processing of low-frequency object information (Bar et al., 2006; Goddard et al., 2016). Third, as subjects’ task was unrelated to object recognition (passive paradigm), it could be concluded that the observed spatiotemporal dynamics of category processing have been hard-wired rather than task-driven. Finally, the results showed that a feed-forward convolutional neural network model could predict the first two suggested stages of visual processing implying that rather than feedback, the second stage of visual processing could be considered a processing pathway operating in parallel to the known processing stages implemented in the ventral visual stream (from V1 to IT).

Although a few recent studies have observed the information regarding the processing of variations in the human brain (Hong et al., 2005; Roth and Zohary, 2015; Goddard et al., 2016; Carlson et al., 2011), the current study is the first to investigate the whole-brain spatiotemporal dynamics of affine (i.e. size and position) and non-affine (i.e. lighting and pose) variation processing using a systematically designed image set. The current results have extended previous studies which concentrated on specific variations (e.g. position; Carlson et al., 2011), evaluated limited areas of the brain (e.g. V4 and IT; Hong et al., 2016), used low temporal resolution recording methods (e.g. fMRI; Roth and Zohary, 2015; Jeong and Xu, 2016) or overlooked possible flows of variation information in the brain (Goddard et al., 2016). Through different analyses (Figs. 4-9), I provided support that significant processing of information about categories and variations was initiated (before 130 ms post-stimulus) in the primary visual cortex and the information was simultaneously sent to peri-frontal areas. This entangled processing of category and variation information was also reported by previous studies which suggested the processing of category and variations in alternating layers of the ventral visual stream (Roth and Zohary, 2015; Hong et al., 2016). In the later window (from 150 to 400 ms), which overlapped with the peri-frontal to peri-occipital window of information flow (Fig. 9A), variation and category information showed spans of in-phase and anti-phase patterns (Fig. 4A, right). The relationship between the category-variation phase patterns correlation (Fig. 4A, right) and the direction of information flow (Fig. 9A, bottom) suggested distinct mechanisms for transferring information from occipital to frontal areas and back to occipital areas. Previous studies have suggested that the information from early visual cortex travels through fast dorsal magnocellular pathway to reach OFC (Bar et al., 2006; Kveraga et al., 2007). The frontal-to-occipital information, however, may use slower pathways which start from frontal cortex and end at occipitotemporal and temporal areas (Peyrin et al., 2010; Chen et al., 2006; Pantazatos et al., 2012; Noudoost and Moore, 2010; Rottschy et al., 2013). The final window (starting from 450 ms), showed simultaneous processing and transferring of category and variation information from occipital to frontal areas. During this final window, the representational information was probably transferred to peri-frontal areas for final cognitive processes such as decision-making and action preparation (Kadohisa et al., 2013; Stokes et al., 2013).

Although several studies have recently proposed the contribution of peri-frontal areas (LOC) to the encoding of low-frequency object information (Bar et al., 2006; Goddard et al., 2016), the current study seems to be the first to show that information regarding variations was also processed by peri-frontal areas. It did not only show that peri-frontal area contributed to variation processing, but also revealed that this area sent variation information to occipital areas. This implies that, in contrast to the consensus that the ventral and dorsal visual streams dominate the processing of category and variation information by feed-forward mechanisms (Reisenhuber and Poggio, 2002; Sereno and Lehky, 2010), the representations in peri-frontal cortex can provide even larger amounts of information compared to those areas in considerable spans of the processing window (Fig. 8, compare the time windows after 200 ms). Results also showed an earlier (from 0 to 50 ms) rise of the information curves in peri-frontal than in peri-occipital areas (Fig. 4B, C and Fig. 8) which can be explained by a previously suggested “framing model” (Chen et al., 2006; Horr et al., 2014). In this model, a frame of the object, which is a vague low-frequency representation or the gist of it, is constructed in peri-frontal brain areas, enhanced by the peri-occipital to peri-frontal flows through magnocellular pathways and is fed back during the second stage of processing to enhance the details of object representations for accurate recognition.

As I did not separate the low‐ and high-frequency components of the objects, it could not be determined which object features were processed by the peri-frontal areas. For variations, however, an advantage was observed in the processing of variations of lighting and pose (rather than variations in size and position) in the peri-frontal areas compared to peri-occipital areas throughout the processing time (Fig. S3B). This is on par with previous suggestions that frontal areas may play role in the compensation of non-affine variations (Karimi-Rouzbahani et al., 2017b; Serre et al., 2005).

An interesting implication of the current results was that the observed role of peri-frontal brain areas in object and variation processing seemed to be hard-wired in the processing procedure rather than activated by top-down cognitive processes which are generally activated during recognition. In fact, as the current paradigm was irrelevant to object recognition, the contribution of the frontal brain could not have resulted from subject’s task. This aspect of the results, while supporting previous findings on the role of peri-frontal cortex in object recognition (Bar et al., 2001; Bar et al., 2006; Goddard et al., 2016), has extended those results to the general case of object processing as opposed to object recognition which can be highly affected by the task, context, etc. (Bugatus et al., 2017; Brab et al., 2013; Karimi-Rouzbahani et al., 2017c). This is on par with a recent study which has suggested that the activation of orbitofrontal cortex is independent of the task, the characteristics of the visual input and explicit recognition of the stimuli (Horr et al., 2014).

The two-stage processing of variations observed in the computational model (Fig. 10, stage one: layers 1-4 and stage two: layers 5-7) can be explained by the dimensionality of the model’s representational space at different model layers. In other words, a very high correlation (r = 0.7117, p = 0.0728, n=7; Pearson linear correlation) was observed between the size of the representational space (i.e. which are 69987, 43264, 64896, 64896, 9216, 4096 and 4096 respectively for layers 1 to 7) and the decoding indices averaged across model layers (Fig. 10A). The correlation was much less for lighting alone (r = 0.4876, p = 0.267) which was probably a result of the lighting, as a low-level image feature, being compensated for in the first layer of the model. Moreover, the category decoding curve and the model’s dimensions showed an anti-correlated pattern (r = ‐0.9231, p = 0.003). It can be concluded that, as suggested by the idea of population coding of variations (Rust and DiCarlo, 2010), large neural populations may be proper candidates to compensate for object variations, while for the encoding of categories, the brain may exploit its inter-layer connections (as supported by the two final fully-connected layers of the computational model (Krizhevsky et al., 2012)).

While some recent studies have reported correlated processing stages between human brain and those computational models (as in Fig. S4; Cichy et al., 2016; Hong et al., 2016) especially at higher visual areas (Khaligh-Razavi and Kriegeskorte, 2014; Cadieu et al., 2014), they have overlooked possible parallel processing mechanisms that could have contributed to those correlated patterns. In other words, computational models and the brain might have implemented different sets of strategies to reach the same abstract representations of objects found in their final processing stages as previously proposed (Karimi-Rouzbahani et al., 2017c). Therefore, even a layer-wise correlation does not rule out the possible existence of parallel mechanisms for object processing in the brain.

One advantage of this study to a most relevant study, which supported the role of peri-frontal areas in object recognition (Goddard et al., 2016), is that it used a rapid presentation paradigm in which objects were presented only for 50 ms. This was important since longer presentation times could have caused the dominance of peri-occipital to peri-frontal information (referred to as feed-forward) flow compared to the peri-frontal to peri-occipital (feedback) flow, leading to the underestimation of feedback influence. As argued by the authors (Goddard et al., 2016), it may explain the dominance of feedforward as well as the lag of feedback flows in that study. However, these results suggest the need for a systematic study to investigate the impact of presentation time on the amplitude and the temporal dynamics of object information flow in the brain.

While the results of current study have provided new insights into the spatiotemporal dynamics of category and variation processing in the human brain, they raised several new questions. Does the task affect the processing of variation information differently from category information? It has been recently shown that the task can significantly modulate category representations in high-level vision-related areas (Bugatus et al., 2017) which can result in the modulation of processing strategies (Karimi-Rouzbahani et al., 2017c). It remains to be studied for variations as well. Second, is the variation information observed here also observed when objects are presented simultaneously in complex scenes? Third, what are exactly the regions which process category and variation information in peri-frontal areas? Although there are suggestions on the role of orbitofrontal cortex in that regard, more accurate recording methods (e.g. combined EEG and fMRI) can tell if information about categories and variations are processed by the same or different brain regions. Regardless of how these questions are addressed, the current investigation provides new insights into the role of peri-frontal brain areas in category and variation processing.

